# Functional Bias of Contractile Control in Mouse Resistance Arteries

**DOI:** 10.1101/2024.04.03.588016

**Authors:** Nadia Haghbin, David M. Richter, Sanjay Kharche, Michelle S.M. Kim, Donald G. Welsh

## Abstract

**Background:** Constrictor agonists set vascular tone through two coupling processes, one tied to (electromechanical), the other independent (pharmacomechanical) of membrane potential (V_M_). This arrangement raises an intriguing query: are they variably recruited such that each agonist elicits a range of vasomotor signatures, functionally biased towards one mechanism or the other? This query underlies this study and our examination of agonist-induced arterial constriction.

**Methods:** Mouse mesenteric arteries were exposed to a classic G_q/11_ (phenylephrine) or G_q/11_/G_12/13_ (U46619) coupled receptor agonist, and responses monitored in the absence and presence of L-type Ca^2+^ channel/protein kinase inhibitors. Contractile work was supplemented with measures of protein phosphorylation, V_M_, and cytosolic Ca^2+^; conceptual insights were enhanced with computational modeling.

**Results:** Each constrictor elicited a response curve that was attenuated and rightward shifted by nifedipine, findings aligned with functional bias; electromechanical coupling preceded pharmacomechanical, the latter’s importance rising with agonist concentration. Ensuing contractile and phosphorylation (CPI-17 & MYPT1 (T-855 & T-697)) measures revealed phenylephrine-induced pharmacomechanical coupling was tied to protein kinase C (PKC), while U46619 was tied to both PKC and Rho-kinase. A switch to pharmacomechanical coupling dominance occurred when agonist superfusion was replaced with discrete application to a small portion of artery. This switch was predicted by electromechanical modeling and supported by direct measures of V_M_ and cytosolic Ca^2+^.

**Conclusions:** Our work illustrates that constrictor agonists elicit functionally biased responses and that arteries toggle among contractile mechanisms, dependent on receptor signal bias, structural/electrical properties, and how agents are applied. We discuss how hemodynamic control is intimately tied to functional bias in both health and disease states, including but not limited to arterial vasospasm.

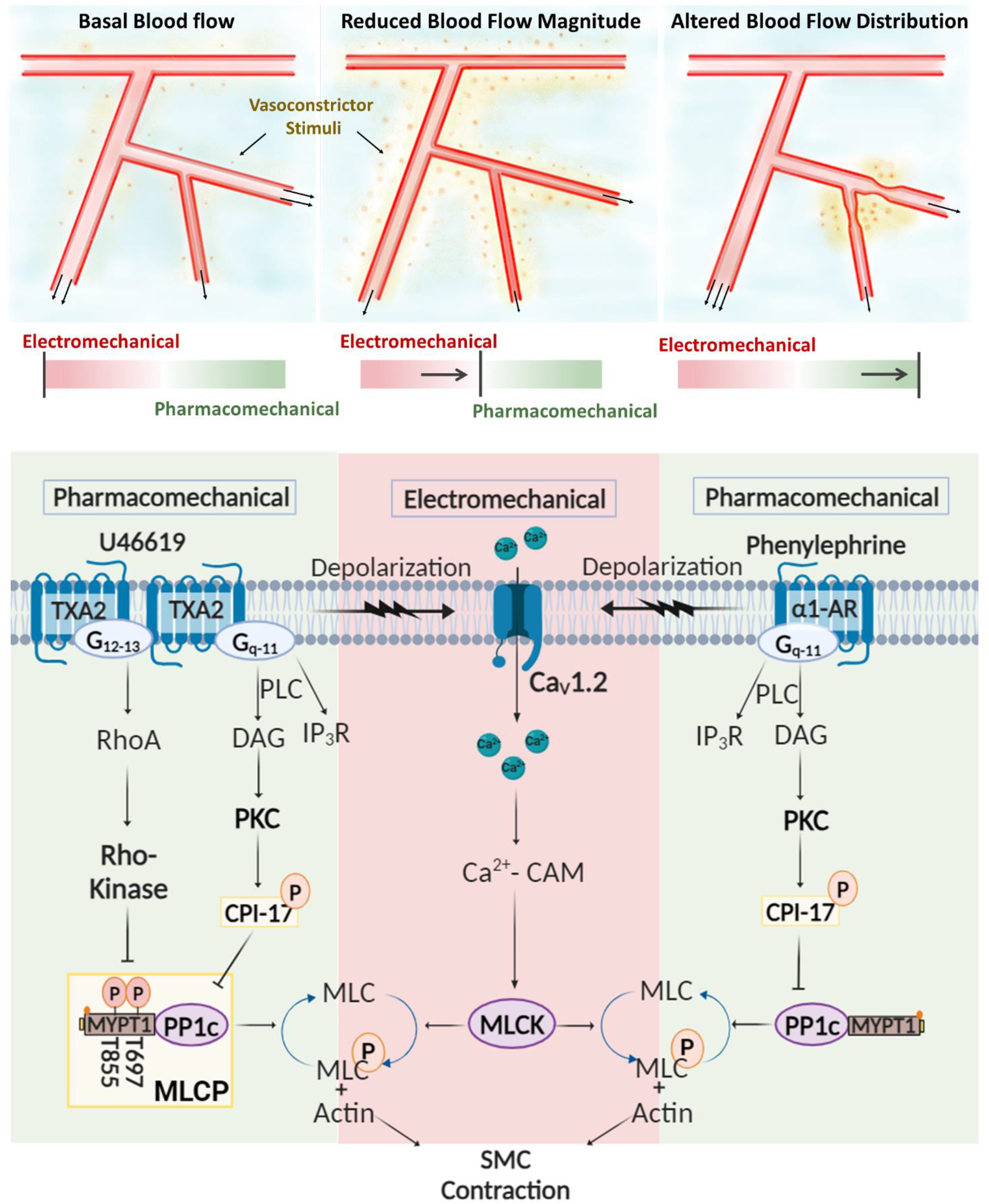

**Highlights:**

- Agonist-induced constrictor responses exhibit a “functional bias” toward electromechanical or pharmacomechanical coupling dependent on agent concentration and mode of application.
- Electromechanical coupling typically but not exclusively precedes pharmacomechanical, the latter rising to prominence with agent concentration. Pharmacomechanical coupling is mediated through receptor pathways linked to PKC and Rho-kinase, regulatory proteins that target the catalytic and targeting subunit of MLCP.
- Functional bias is malleable and thus each agonist elicits a range of vasomotor signatures, each presumptively important in controlling blood flow delivery in space and time.

## Introduction

Resistance arteries form an integrated vascular network responsible for optimizing blood flow delivery to metabolically active cells^1,2^. In peripheral tissue, like skeletal muscle or the gut, basal tone is initially set by intravascular pressure and the release of neurotransmitters from sympathetic nerves. Each constrictor stimuli works through G-protein coupled receptors and associated transduction pathways to govern electromechanical and pharmacomechanical coupling, processes that set the phosphorylation state of the myosin light chain^3,4,5^. Electromechanical coupling is defined as the modulation of myosin light chain kinase (MLCK) through changes in membrane potential (V_M_) that drive an influx of Ca^2+^ through voltage-gated Ca^2+^ channels^6,7,8^. Pharmacomechanical coupling classically refers to the regulation of myosin light chain phosphatase (MLCP) through signal pathways that activate two key kinases, the first protein kinase C (PKC), the second Rho-kinase. The former targets CPI-17 (myosin phosphatase inhibitory protein, 17 kDa), a modulator of the MLCP catalytic subunit, PP1cδ, whereas the latter phosphorylates MYPT1, the targeting subunit of MLCP, via its T-697 and T-855 regulatory sites^9,10,11^.

Vascular studies have long recognized that constrictor agonists use both electro- and pharmacomechanical coupling to set arterial tone^12^. This knowledge raises a fundamental question: why do agonists engage two distinct coupling mechanisms when one is seemingly sufficient to elicit constriction? One answer conceivably lies in deepening our understanding of blood flow control and recognizing that each vessel segment needs to engage in a breadth of vasomotor behavior (spatial and temporal) to optimize the magnitude and distribution of delivery in real time^13^. This necessity is particularly clear in computer modeling which has noted blood flow optimization is near impossible in networks encoded with just one coupling process, for example electromechanical control^14,15^. Consequently, a second contractile process, one mechanistically distinct and variably engaged relative to the first would be seemingly important^16^. The idea that the coupling mechanisms aren’t fixed in proportion but rather “functionally biased” toward electro- or pharmacomechanical coupling is a novel concept worthy of further interrogation.

This study determined whether functionally biased responses were observable in mesenteric arteries exposed to a G_q/11_ (phenylephrine) or G_q/11_/G_12/13_ (U46619) coupled receptor agonist, analogs akin to those released from perivascular nerves or endothelial cells/platelets, respectively.

Our examination began by monitoring concentration response curves prior to and following pharmacological manipulation and subsequently to measurements of protein phosphorylation, membrane potential (V_M_), and cytosolic Ca^2+^ to garner deeper insights. We observed compelling evidence of functional bias, each agonist eliciting a contractile response where electromechanical coupling preceded pharmacomechanical, the latter increasing in prominence as concentrations rose. The pharmacomechanical coupling response induced by phenylephrine was strongly linked to PKC, whereas that induced by U46619 was tied to both PKC and Rho-kinase activation. Ensuing experiments demonstrated how functional bias switches to complete pharmacomechanical dominance, by restricting the agent’s application to small portions of the artery, a finding predicted from computational modeling. We conclude from this foundational work that constrictors elicit functionally biased responses and that arteries toggle between the contractile mechanisms. We discuss how this knowledge advances our mechanistic understanding of hemodynamic control in health and disease states, including arterial vasospasm.

## Methods

### Animal and tissue preparation

Animal procedures were approved by the animal care committee at the University of Western Ontario, in accordance with guidelines set forth by the Canadian Council on Animal Care and ARRIVE guidelines. Male C57BL/6J mice (wild type, 16-20 weeks of age) obtained from Jackson Laboratories were euthanized by CO_2_ asphyxiation^17^. The mesentery carefully removed and placed in cold PBS solution (pH 7.4) containing (in mM): 138 NaCl, 3 KCl, 10 Na_2_HPO_4_, 2 NaH_2_PO_4_, 5 glucose, 0.1 CaCl_2_, and 0.1 MgSO_4_. Fourth order arteries were dissected free of connective tissue and cut into 2 – 3 mm segments for further experimentation^18^. Mesenteric arteries from male Ca_V_3.1^-/-^ mice (global knockout, in house colony; 16-20 weeks of age) were also used in one experimental subset; homozygotes were generated from Ca_V_3.1^-/-^ mice crossed onto a C57BL/6J background^19^.

### Vessel Myography

Isolated mesenteric arteries were placed in a pressure myograph system, cannulated, and equilibrated (intravascular pressure, 15 mmHg for 20 min) with warm (37°C) physiological saline solution (PSS; 5% CO2, balance air) containing (in mM): 119 NaCl, 4.7 KCl, 1.7 KH_2_PO_4_, 1.2 MgSO_4_, 1.6 CaCl_2_, 10 glucose, and 20 NaHCO_3_. Vessel reactivity was then assessed by applying 60 mM KCl to the bath and measuring the diameter via an automated edge detection system (IonOptix, MA) and a 10x objective. Following wash off in standard PSS, intravascular pressure was incrementally increased to 60 mmHg and two sets of experiments were performed. First, we assessed the responsiveness of mesenteric arteries to superfused phenylephrine (3x10^-9^ to 10^-4^ M) or U46619 (10^-9^ to 10 ^-6^ M) in the absence and presence of YM254890 (0.1 µM, G_q-11_ inhibitor), nifedipine (0.3 µM, L-type Ca^2+^ channel blocker), calphostin C (0.3 µM, PKC inhibitor), or Y-27632 (20 µM, Rho-kinase inhibitor). The percent maximal constriction was calculated as [*100×(D0–D)/D0-Dm*], where *D0* is diameter at 60 mmHg (no agonist), *D* is the diameter at each agonist concentration, and the *Dm* is the maximal constriction at the highest agonist concentration under control conditions. In the second experimental set, phenylephrine (100 µM) and U46619 (10 µM) were focally applied upstream through small-bore glass micropipettes (1-2 μm tip, pressure ejection (30 psi, 10s pulse)), in the absence and presence of nifedipine (0.3 µM) or calphostin C (0.3 µM); diameter was measured proximal to and distal from the site of agonist application.

### Arterial membrane potential (V_M_)

Arterial V_M_ was assessed by inserting a glass microelectrode backfilled with 1 M KCl (tip resistance = 90-110 MΩ) into the arterial wall^20^. V_M_ was first assessed under control conditions and following global application of phenylephrine (30 µM) or U46619 (0.1 µM) in the superfusate. In a second set of experiments, V_M_ was measured prior to and following the focal stimulation by phenylephrine (100 µM, 10s pulses) or U46619 (10 µM, 10s pulses). Criteria for a successful recording are (1) sharp negative V_M_ deflection on electrode insertion; (2) stable V_M_ reading for a minimum of 2 minutes after entry; and (3) sharp return to baseline on electrode removal. To prevent vessels from overly constricting and introducing a motion artifact, 20 µM Y-27632 was present in the superfusate^21^.

### Intracellular Ca^2+^ ([Ca^2+^]_i_) Measurements

To evaluate [Ca^2+^]_i_, mouse mesenteric arteries were isolated and loaded with 70 µM Fura-2-AM (dissolved and diluted in 50µl DMSO, 3.4 µl Pluronic acid and 955 µl HBSS buffer) for 1 hour at room temperature in the dark^22^. Arteries were then mounted in a pressure myograph and placed on top of an inverted epifluorescence microscope. Fura-2 was alternatingly excited at 340 nm and 380 nm (200 ms alternating intervals); emission spectra (510 nm) was collected on a RETRA light engine camera (Lumencor) viewed through a 10X objective (1.2 NA). Data was analysed using Nikon NIS Elements (AR 4.20.01) software. Background fluorescence was subtracted and the ratio of emitted fluorescence (F_340_/F_380_) calculated as a measure of [Ca^2+^]_i_. Experimentally, [Ca^2+^]_i_ was continuously monitored while vessels were treated with phenylephrine (100 µM focal or 30 µM global) or U46619 (10 µM focal or 0.1 µM global) in the presence or absence of diltiazem (30 µM, an L-type Ca^2+^ channel blocker with reduced photosensitivity). [Ca^2+^]_i_ was monitored at baseline and once vasomotor tone had peaked to agonist application.

### Western blot analysis

A three-step western blotting protocol was used to quantify the phosphorylation state of MYPT1 (T-855 or T-699) and CPI-17 to agonist application^23^. Mesenteric arteries were cut into 2 mm lengths, with two segments placed together in a conical vial containing Ca^2+^ PSS (37°C, 5% CO_2_, balanced air; 15 min). After equilibration, one vial was left untreated and a second exposed to phenylephrine (30 µM) or U46619 (0.1 µM) for 3 minutes. Immediate fixation in buffered acetone (10% trichloro acidic acid plus 10 mM dithiothreitol in acetone) was followed by freeze drying (- 55 °C, overnight). Freeze dried tissues were then transferred to a sample buffer (75 mM Tris HCL (pH 6.8), 10% Glycerol, 6% SDS, Bromophenol Blue, 2-mercaptoethanol) and vortexed vigorously. Following extraction, proteins were separated on a 4% to 10% gradient SDS-PAGE or a phosphate-affinity tag SDS-PAGE (CPI-17), and then were transferred to nitrocellulose membranes^24^. Membranes were blocked for 1 hour (5% nonfat dairy milk in Tris-buffered saline), and then incubated with a primary antibody (p-MYPT1-T697, p-MYPT1-T855, CPI-17; 1:500) followed by a biotin-conjugated secondary antibody (1:5,000) and HRP-conjugated streptavidin (1:10,000). The membrane was washed repeatedly with Tris Buffered Saline with Tween (TBST) after each incubation step. The blot was developed using Amersham ECL Prime Western Blotting Reagent and imaged on a Gel Doc using Image Lab software (Bio-Rad). The density of each band was quantified using scanning densitometry^24^. MYPT1 phosphorylation was normalized to α- smooth muscle actin; phosphorylated CPI-17 was normalized to total CPI-17 expression. Agonist treated vessels were then normalized in relation to untreated vessels.

### Computational modeling Construction of virtual artery

A 500 µm bilayer artery model with an inner endothelial cell (EC) layer and a surrounding smooth muscle cell (SMC) layer was developed (refer to Figure 6A)^14,25^. Cells were electrically coupled with gap junctional resistance set at 3 MΩ (EC-to-EC), 90 MΩ (SMC-SMC), and 1800 MΩ (EC-SMC; note each smooth muscle cell was linked to two ECs. Reaction currents simulated ion channel activity; arterial diameter was initially set at 75 µm, and was adjusted to SMC membrane potential changes, as modeled through ordinary differential equations^14,26^. Simulations consisted of injecting a variable number of smooth muscle cells (All, 1/2, 1/4, 1/8) with a current of +0.83 pA. The depolarizing current when applied to all smooth muscle cells elicited a +12 mV depolarization, a response in line with physiological observations^27^. Greater detail as to model construction can be found in the original publication^14^.

### Numerical methods

A Mersenne Twister algorithm facilitated random EC-SMC coupling, with cell electrical activity computed by our ordinary differential equations using a high-order implicit backward difference formula^28,29^. Distributed memory parallelization was implemented using the MPI library^30^. The simulations were advanced over several iterations, until the relative solution changes over consecutive iterations were negligible (i.e., relative tolerance less than 10^-6^) at which point steady state was assumed. The simulations were performed on a local HPC cluster running Fedora 31.

### Statistical analysis

Data are expressed as mean ± SEM and n indicates the number of samples or arteries. Nonlinear curve fitting method of GraphPad Prism 10.1 software was used to analyze the accumulative dose-effect curves. Two-way ANOVA and *t* tests (paired or unpaired) were performed to compare the effect of a given condition/treatment on diameter, V_M_, intracellular [Ca^2+^], or phosphorylation state. *P* values ≤0.05 were considered statistically significant.

### Solution and chemicals

Primary and secondary antibodies were obtained from the following sources: Anti-Phospho-MYPT1 (Thr697) and (Thr855) rabbit polyclonal antibody was purchased from New England Labs. Anti-α smooth muscle actin rabbit polyclonal antibody was purchased from Abcam. Anti-CPI-17 antibody rabbit polyclonal IgG, and Biotin-sp conjugate goat anti-rabbit IgG antibody Millipore Sigma. Y-27632 and U46619 were purchased from Tocris. YM254890 was acquired from Focus Biomolecules and calphostin C was acquired from Cayman Chemical. Nifedipine, phenylephrine hydrocholoride, diltiazem hydrocholoride, Streptavidin-peroxidase and Fura-2 LR/AM and all other chemicals were obtained from Millipore Sigma, unless stated otherwise.

Where DMSO was used as a solvent, the maximal DMSO concentration after application did not exceed 0.05%.

## Results

### G-Protein coupling and the *α*-1 Adrenergic and Thromboxane A2 (TXA_2_) receptors

To probe the nature of G-protein coupling, agonist-response curves to phenylephrine or U46619 were constructed in the absence and presence of a G_q/11_ inhibitor (YM254890)^31,32^. Figure 1 reveals that phenylephrine (3x10^-9^ M to 10^-4^ M) and U46619 (10^-9^ M to 10^-6^ M) induced robust concentration-dependent constrictions in mouse mesenteric arteries. Superfusion of YM254890 abolished phenylephrine-induced constriction (Figure 1A) while its impacts on the U46619 response was mixed; a near complete elimination was observed when agonist concentration was low but only partial at higher concentrations (Figure 1B). These findings align with α1-adrenoreceptors being singularly coupled to G_q/11_ while that of the TXA_2_ receptor was tied to both G_q/11_ and G_12/13_.

**Figure 1:**
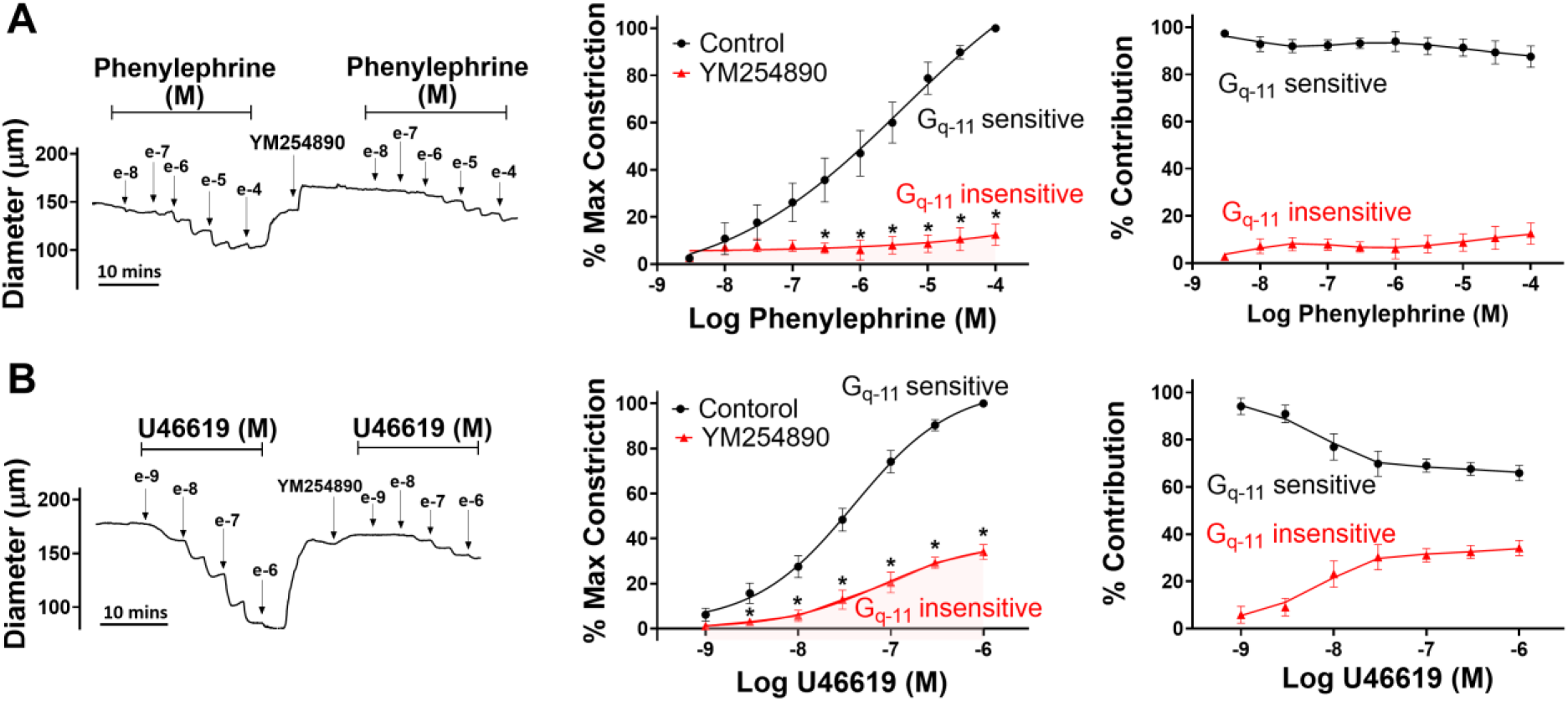
Effect of G_q-11_ inhibitor on agonist induced constriction. Isolated mesenteric arteries from C57BL/6 mice were exposed to phenylephrine or U46619 in presence and absence of 0.1 µM YM254890 (G_q-11_ inhibitor). **A, B)** Representative traces and summary data (% maximum constriction; % contribution of G_q-11_ sensitive and insensitive components) were calculated. N= 6 arteries from 6 mice for each experiment. Data are mean ± SEM. * indicates a significance vs control (Two-way ANOVA, P < 0.05)

### Sequential activation of contractile mechanisms

To address how electro- and pharmacomechanical coupling are engaged, agonist-response curves were constructed in the absence and presence of nifedipine, an L-type Ca^2+^ channel blocker that uncouples V_M_ from arterial constrictor control. Akin to the preceding work, the superfusion of phenylephrine (3x10^-9^ M to 10^-4^ M) or U46619 (10^-9^ M to 10^-6^ M) induced constriction of mouse mesenteric arteries in a concentration dependent manner (Figure 2A & 2B, left and middle). The subsequent bath application of nifedipine diminished vessel reactivity and shifted the concentration-response curves to both agonists rightward. The nifedipine-sensitive component (electromechanical) dominated tone control at low agonist concentrations whereas the nifedipine-insensitive component (pharmacomechanical) rose to prominence at higher concentrations (Figure 2A & 2B, right). Such findings are indicative of a functional bias in vascular reactivity, with electromechanical coupling preceding pharmacomechanical. Intriguingly, nifedipine’s impact on phenylephrine-induced constriction was more pronounced than on U46619, the latter transduced through the TXA2 receptor via G_q/11_ and G_12/13_ (Figure 1A & 1B, right). Deeper analysis revealed that time-to-peak constriction (phenylephrine), was also moderated by nifedipine application, a finding in line with quicker temporal engagement of electromechanical coupling; a similar time shift wasn’t observed with U46619 (Figure 2C & 2D). These time-dependent assessments were conducted at agonist concentrations that elicited approximate half-maximal constriction.

**Figure 2:**
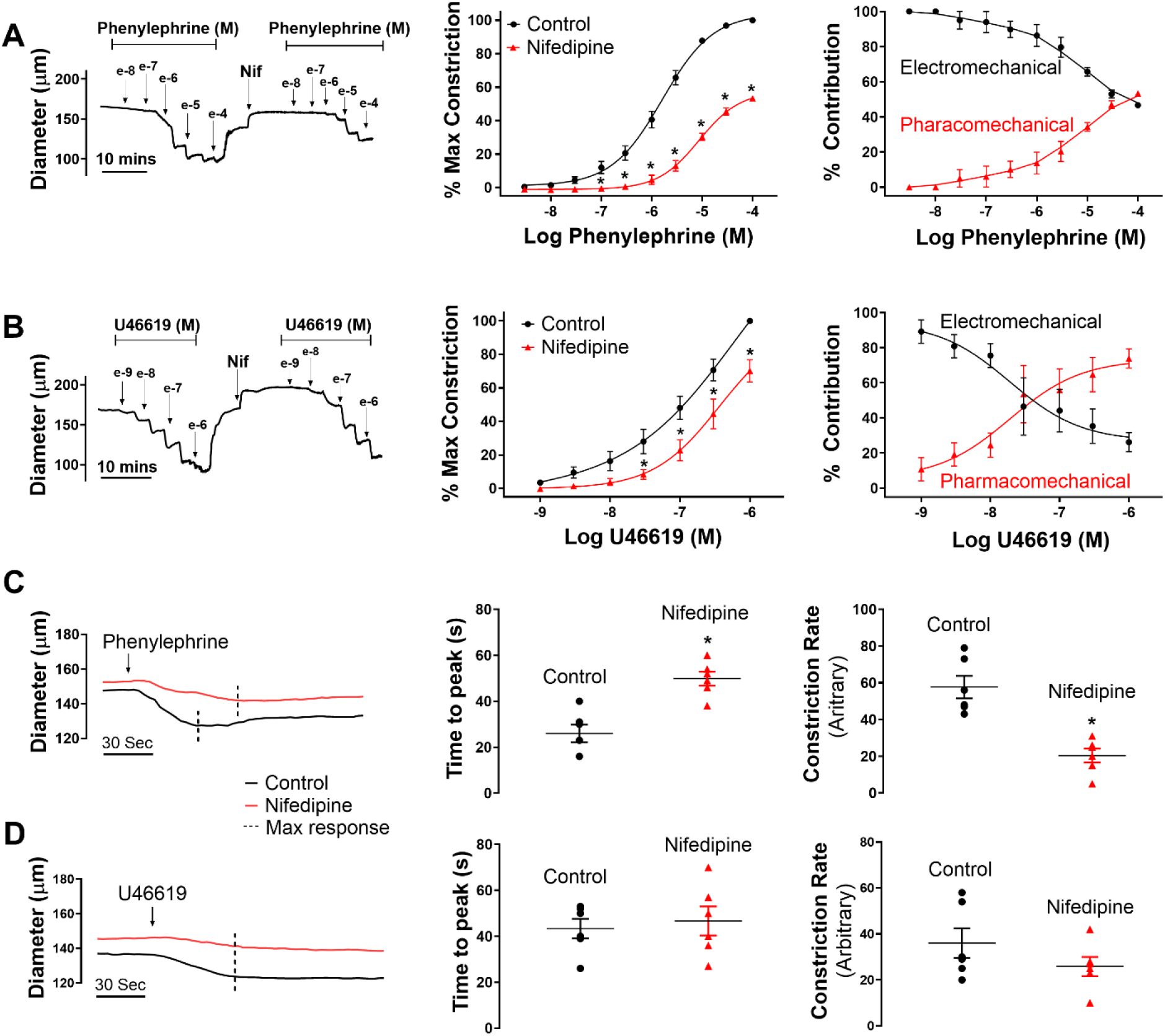
Relative contribution of electro- and pharmacomechanical coupling to agonist induced constriction. Isolated mesenteric arteries from C57BL/6 (wild type) mice were exposed to phenylephrine (**A**) or U46619 (**B**) in presence and absence of 0.3 µM nifedipine (L-type Ca^2+^ channel blocker). Representative traces and summary data (% maximum constriction; % contribution pharmacomechanical and electromechanical components) were calculated. Time-to-peak constriction to 30 µM phenylephrine **(C)** and 0.1 µM U46619 **(D)** in isolated mouse mesenteric arteries. N= 6-8 arteries per group, 1 experiment per mouse. Data are mean ± SEM. * indicates a significance vs control (Two-way ANOVA, paired *t*-test, P-value < 0.05)

In recognition that T-type Ca_V_3.1 channels are expressed in vascular smooth muscle and could confound the interpretation of Figure 2A & 2B, experiments were repeated in mesenteric arteries from Ca_V_3.1^-/-^ mice^33,34^. Agonist induced constriction (Figure 3A & 3B, left and middle) in Ca_V_3.1^-/-^ arteries, appeared quantitatively similar to wild type control (Figure 2A & 2B), in the absence and presence of nifedipine to block L-type Ca^2+^ channels. Each agonist increased tone in a concentration dependent manner, with nifedipine reducing and rightward shifting the constrictor response curves. Similar to wild type arteries, electromechanical coupling dominated at low agonist concentrations whereas pharmacomechanical coupling rose in prominence at higher concentrations (Figure 3A & 3B, right).

**Figure 3:**
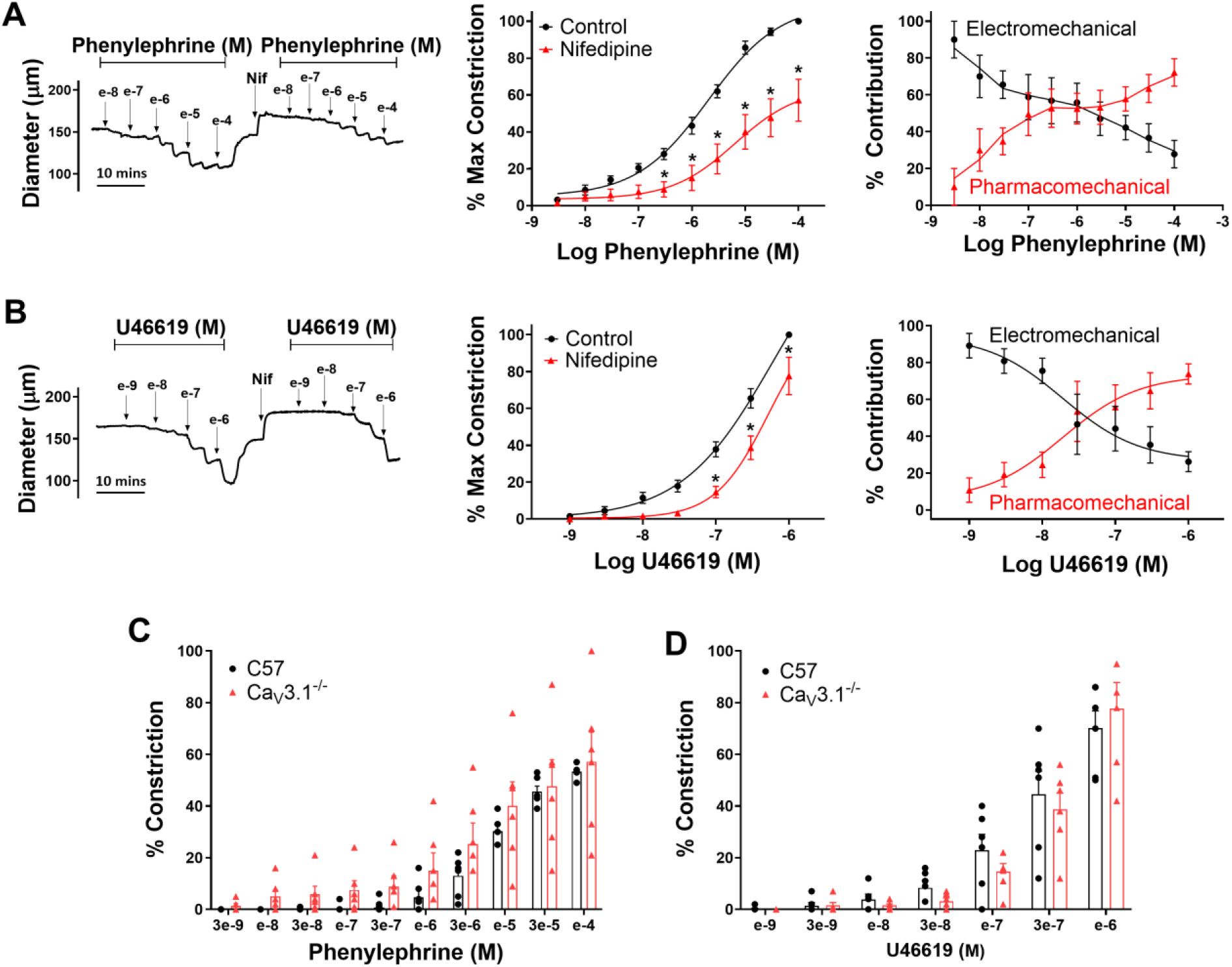
Contribution of T-type Ca^2+^ channels in agonist induced constriction. Isolated mesenteric arteries from Ca_V_3.1^-/-^ mice were exposed to phenylephrine (**A**) or U46619 (**B**) in the presence and absence of 0.3 µM nifedipine (L-type Ca^2+^ channel blocker). Representative traces and summary data (% maximum constriction; % contribution pharmacomechanical and electromechanical components) were calculated. **(E & F)** Summary data compares the % of nifedipine insensitive constriction to phenylephrine and U46619 in arteries from wild type and Ca_V_3.1^-/-^ mice. N= 6-8 arteries per group, 1 experiment per mouse. Data are mean ± SEM. * indicates a significance vs control (Two-way ANOVA, P-value < 0.05)

### Pharmacomechanical coupling and phosphorylation control of MLCP

Pharmacomechanical coupling is intimately tied to MLCP regulation and the phosphorylation of three key sites one on CPI-17, an upstream modulator of PP1cδ, and two on MYPT1 (T-855 & T-697)^35,36^. We employed western blot analysis to examine these sites, prior to and following the application of agonists at concentrations that elicit near maximal constriction. Our findings reveal that a 3 min application of phenylephrine enhanced CPI-17 phosphorylation while having no significant effect on the phosphorylation state of MYPT1 (Figure 4). U46619 application also enhanced CPI-17 phosphorylation, but in addition, markedly elevated MYPT1 T-855 & T-697 phosphorylation. CPI-17 phosphorylation is driven by PKC, and consistent with this kinase driving pharmacomechanical coupling, calphostin C (a pan-PKC inhibitor) fully and partially abolished the nifedipine-insensitive constriction by phenylephrine and U46619, respectively (Figure 5)^11^. Complete elimination of U46619-induced pharmacomechanical response required the further inhibition of Y-27632, a Rho-kinase inhibitor (Figure 5B)^24,37^.

**Figure 4:**
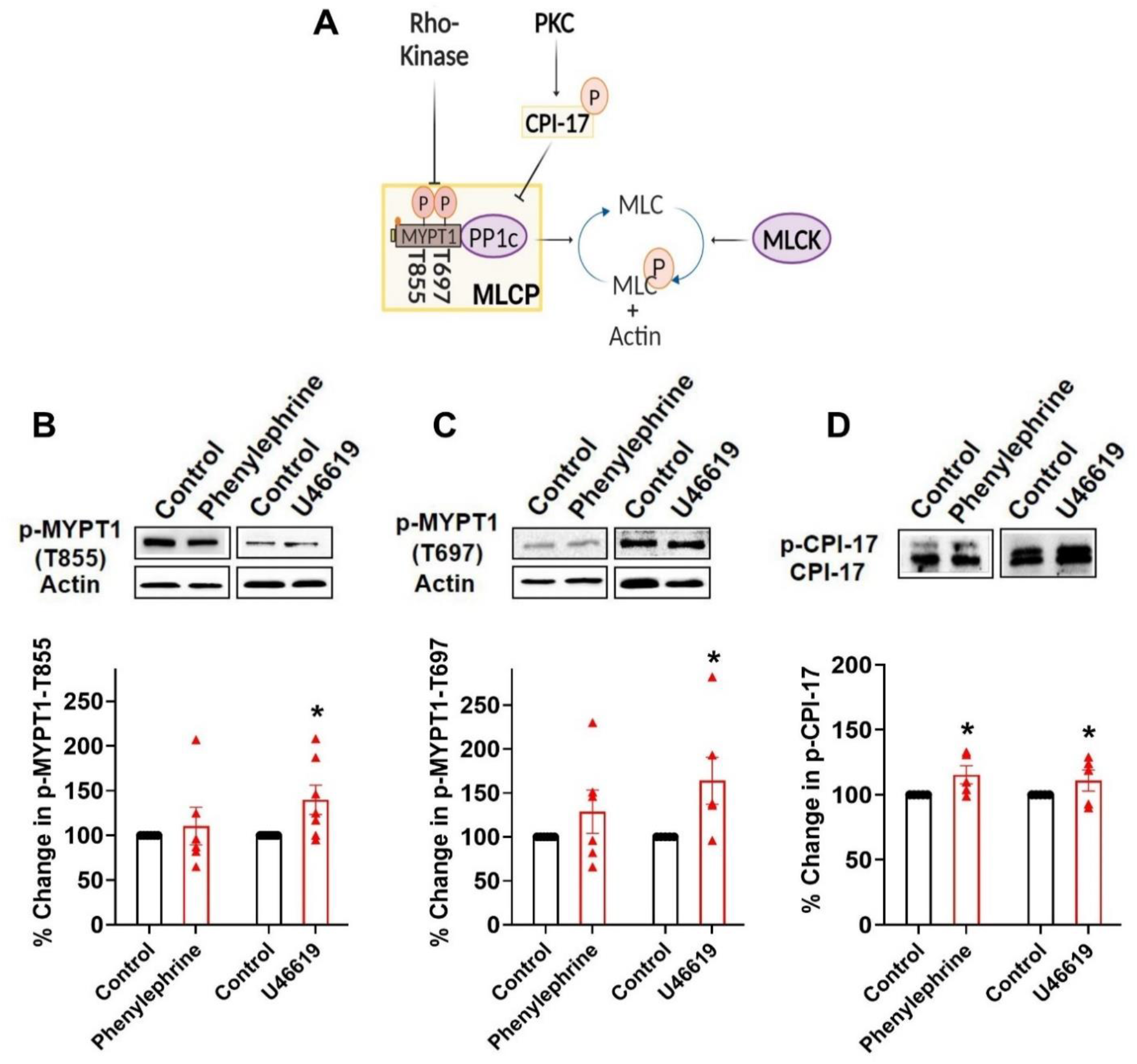
Agonist induced phosphorylation of MLCP regulatory targets. **A)** Key phosphorylation targets (MYPT1, T-855 & T-697 and CPI-17) in response to phenylephrine (30 µM) and U46619 (0.1 µM) application. Summary data shows Phenylephrine treatment increased the phosphorylation state of CPI-17 (**B)**; in comparison U46619 treatment increased the phosphorylation state of CPI-17 (**B**) and MYPT1 (T-855 & T-697) **(C & D)**. α-smooth muscle actin was used as a loading control. N = 3-5 experiments per group, 1 experiment per mouse. Data are mean ± SEM. * indicates a significance vs control (Paired *t-*test, P-value < 0.05)

**Figure 5:**
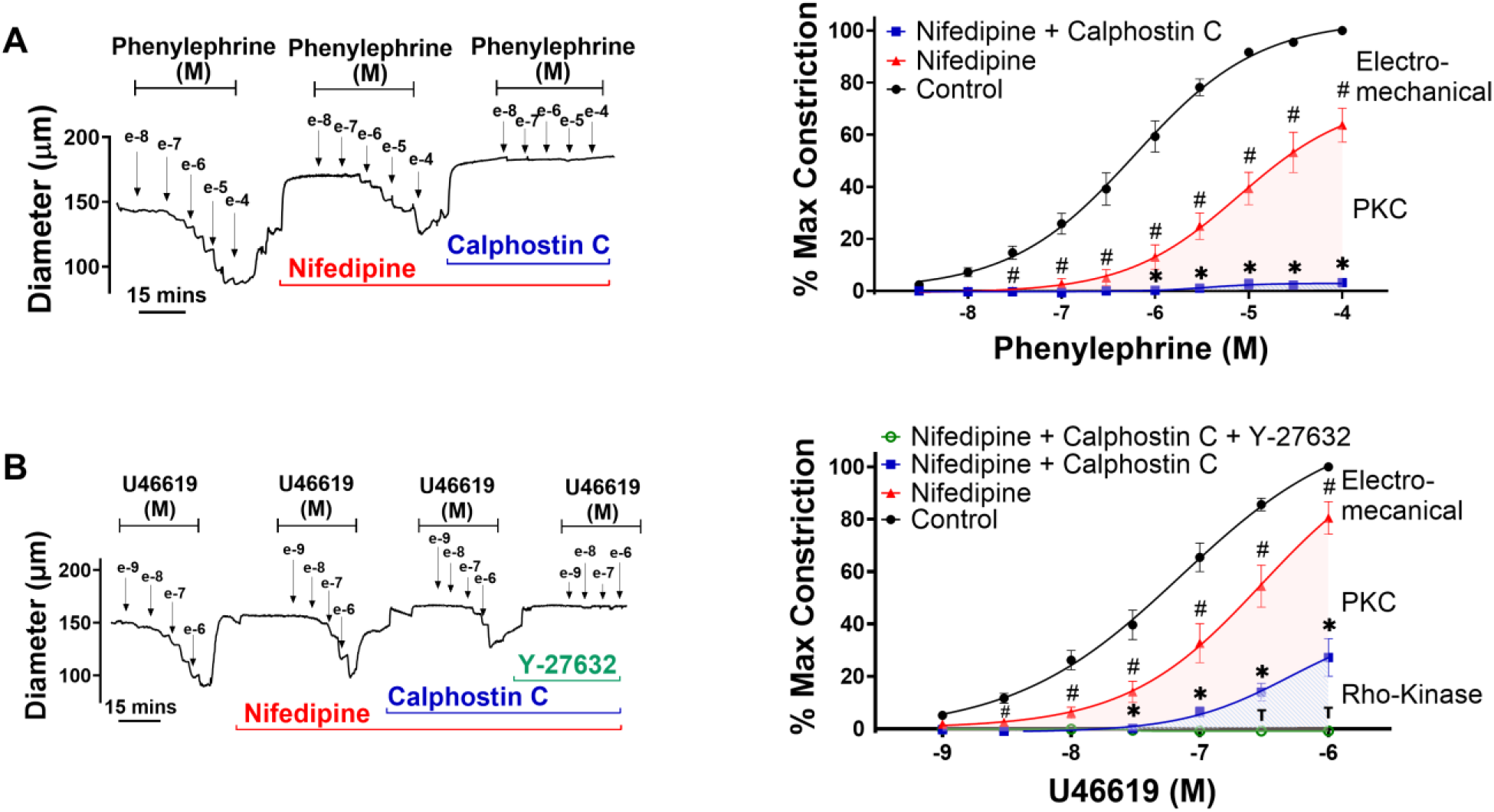
Effects of PKC and Rho-kinase inhibition on agonist induced pharmacomechanical coupling. **A)** Representative trace and summary data of phenylephrine induced constriction with or without 0.3 µM nifedipine and 0.3 µM calphostin C. **B)** Representative trace and summary data of U46619 induced constriction with and without 0.3 µM nifedipine, 0.3 µM calphostin C and 20 µM Y-27632. N=6 arteries per group, 1 experiment per mouse. Data are mean ± SEM. # denotes significant decrease from control curve. * denotes significant decrease from nifedipine-insensitive curve. Ʈ denotes significant decrease from PKC-insensitive curve. Two-way AVOVA, P-value < 0.05.

**Figure 6:**
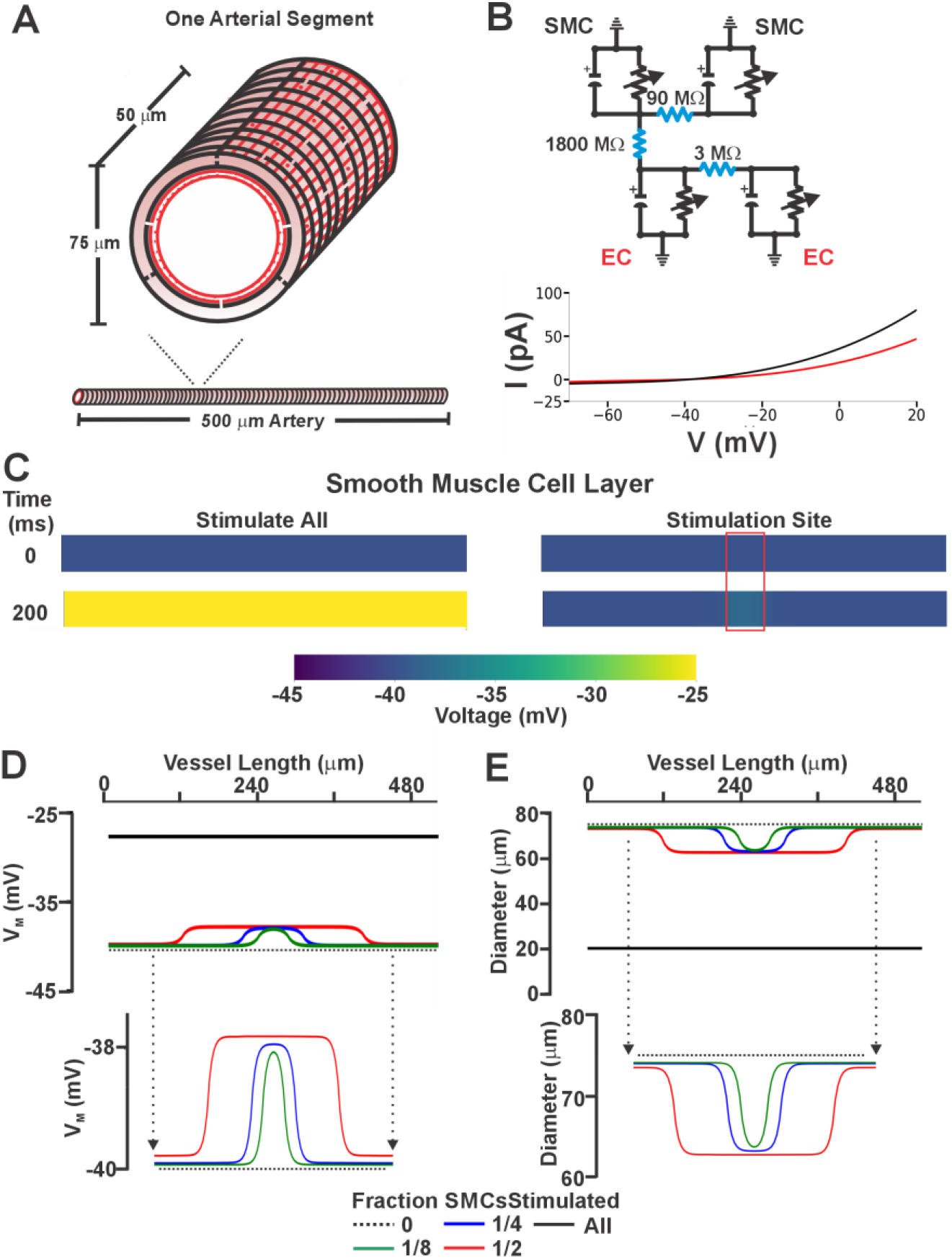
Virtual artery simulations: Smooth muscle cell depolarization in response to current injection. **A)** The virtual artery, measuring 500 µm in length, was composed of a single endothelial layer (depicted in red) and an outer smooth muscle layer (depicted in black). Each section of the artery consisted of 48 endothelial cells and 10 smooth muscle cells. Adjacent smooth muscle cells were electrically connected as were neighboring endothelial cells; each smooth muscle cell was in turn coupled to two endothelial cells. Note in **(B)**, each cell was modeled as a capacitor in parallel with a nonlinear resistor representing the ionic conductivity of the cell membrane. Gap junctions were represented as ohmic resistors. The current-voltage (I-V) characteristics of the nonlinear resistors are shown in the lower panel, with reference to a resting membrane potential of -40 mV. **C)** Color voltage maps of V_M_ as all (left) and 1/8^th^ (right, red box) of all smooth muscle cells were injected with 0.83 pA of depolarizing current (200 ms). (**D &E)** Voltage and vasomotor outcomes for virtual arteries where a 0.83 pA stimulus was injected into all, or 1/2, 1/4 and 1/8 of all smooth muscle cells. Note the stimulation of all smooth muscle cells resulted in the virtual artery depolarized from -40 to –28 mV; stimulation of 1/8 of all smooth muscle cells reduced the electrical response to < 2.0 mV.

### Altering the order and dominance of coupling mechanisms

Given the preceding work, we next considered whether there existed a functional scenario where the order and dominance of coupling mechanisms is reversed. Modeling of electromechanical control reveals one possibility which centers on changing the number of smooth muscle cells generating a depolarizing current^27^. When all smooth muscle cells produce a small depolarizing current, we observed a +12.5 mV response with corresponding constriction, a virtual result that aligns with our experiments in Figure 6. That depolarization, however, decreased markedly to <2 mV as the number of stimulated SMCs was reduced to ^1^⁄_2_ , ^1^⁄_4_ and ^1^⁄_8_ of total. With results implying pharmacomechanical coupling dominates when agonist presentation is restricted, we began testing this scenario by focally applying phenylephrine (100 µM) or U46619 (10 µM) via pipet onto a small portion of a mesenteric artery. Consistent with theory, discrete agonist application elicited a focal constriction resistant to nifedipine, a response indicative of its voltage insensitivity. Further contractile work indicated that pharmacomechanical coupling dominates as PKC blockade (Calphostin C) attenuated the focal constriction enabled by phenylephrine or U46619 (Figure 7). Subsequent V_M_ measurements align with the vasomotor observations, with superfused agonist (phenylephrine or U46619) eliciting a profound arterial depolarization, whereas focal application did not (Figure 8). Note, V_M_ measures were conducted in the presence of Y-27632 to ensure that constriction didn’t introduce a motion artifact into electrical recordings^27^. Furthermore, measures of [Ca^2+^]_i_ supported the previous work in showing that superfused agonist (phenylephrine or U46619) elicited a rise in smooth muscle that was diltiazem (L-type Ca^2+^ channel blocker) sensitive. In contrast, the [Ca^2+^]_i_ response to focal agonist application was markedly muted and resistant to diltiazem, indicating a shift towards pharmacomechanical coupling and the use of a voltage-insensitive Ca^2+^ source (Figure 9).

**Figure 7:**
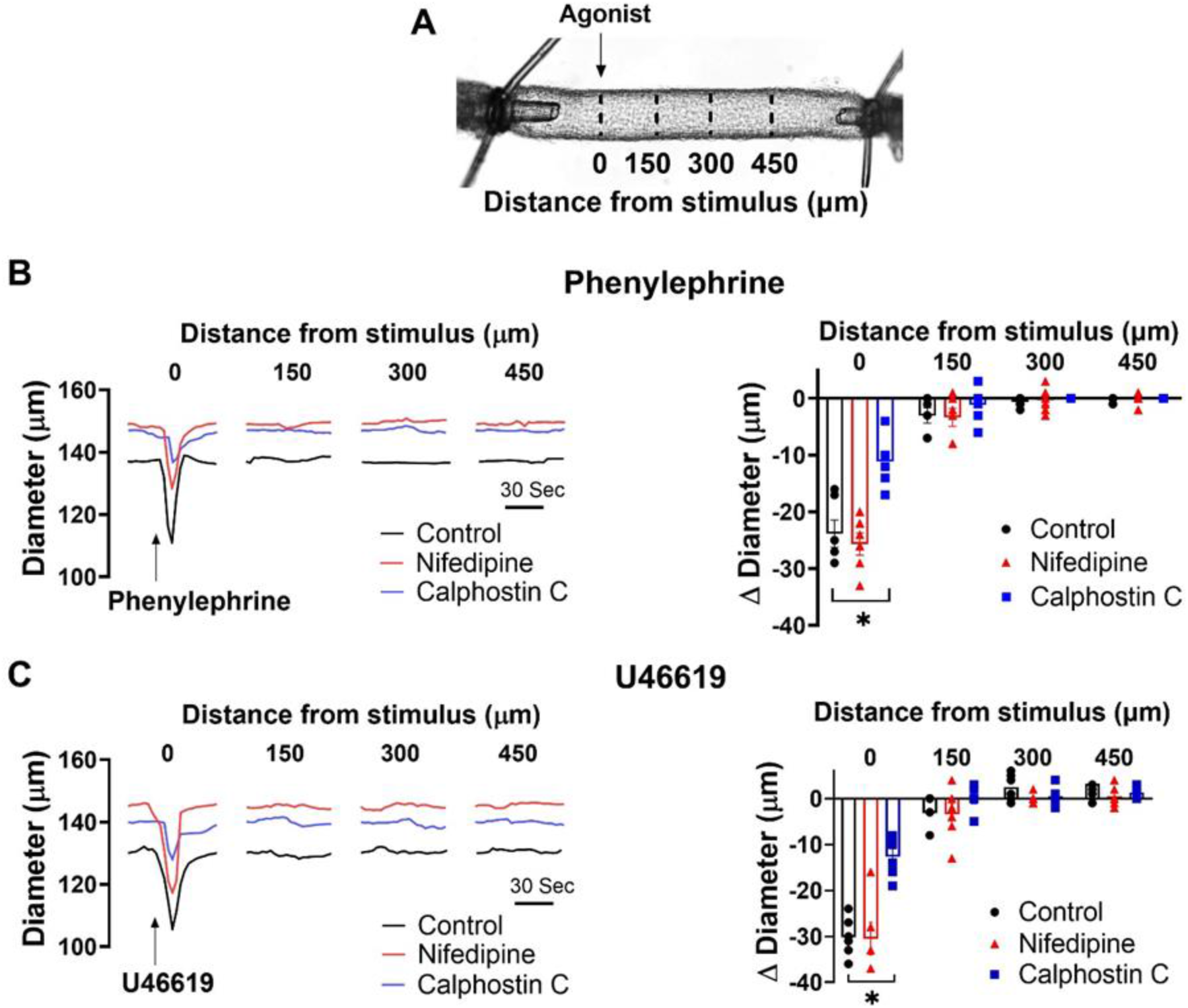
Effects of nifedipine and PKC inhibition on focal constriction. **A)** Agonists were applied onto a small segment of mesenteric arteries using a glass micropipette; diameter was monitored 0–450 μm distal to the point of agent application. Representative traces and summary data of phenylephrine (**B;** 100 µM focally, 10 sec pulse) or U46619 (**C;** 10 µM, 10 s pulse) application, with and without 0.3 µM nifedipine and 0.3 µM calphostin C. N=6 arteries per group, 1 experiment per mouse. * denotes significant decrease vs control (Paired *t* test, P-value < 0.05).

**Figure 8:**
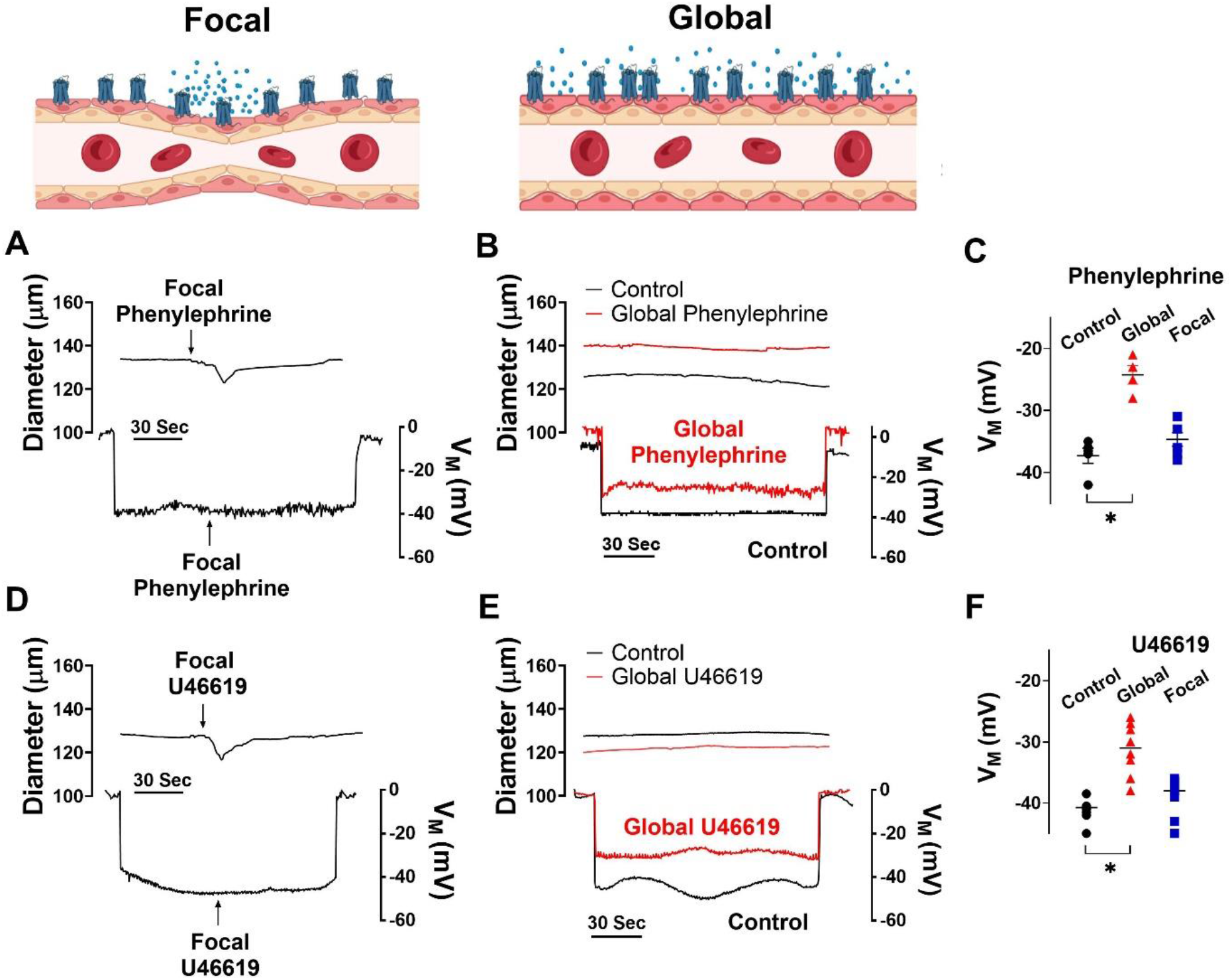
Focal agonist application does not induce arterial depolarization. Isolated mesenteric arteries from C57BL/6 mice were exposed to U46619 (10 µM focally, 10 s pulses or 0.1 μM globally) and phenylephrine (100 µM focally, 10 s pulses or 30 μM globally) was applied to mesenteric arteries while diameter and V_M_ were monitored. Representative tracing **(A & B)** and summary data **(C)** of responses to focal and global phenylephrine. Representative tracing **(D & E)** and summary data **(F)** of responses to focal and global U46619. N = 6 - 8 recording per group; 1 experiment per mouse. Data are mean ± SEM. * denotes significant decrease from control recording (Paired and unpaired *t-*test, P-value < 0.05).

**Figure 9:**
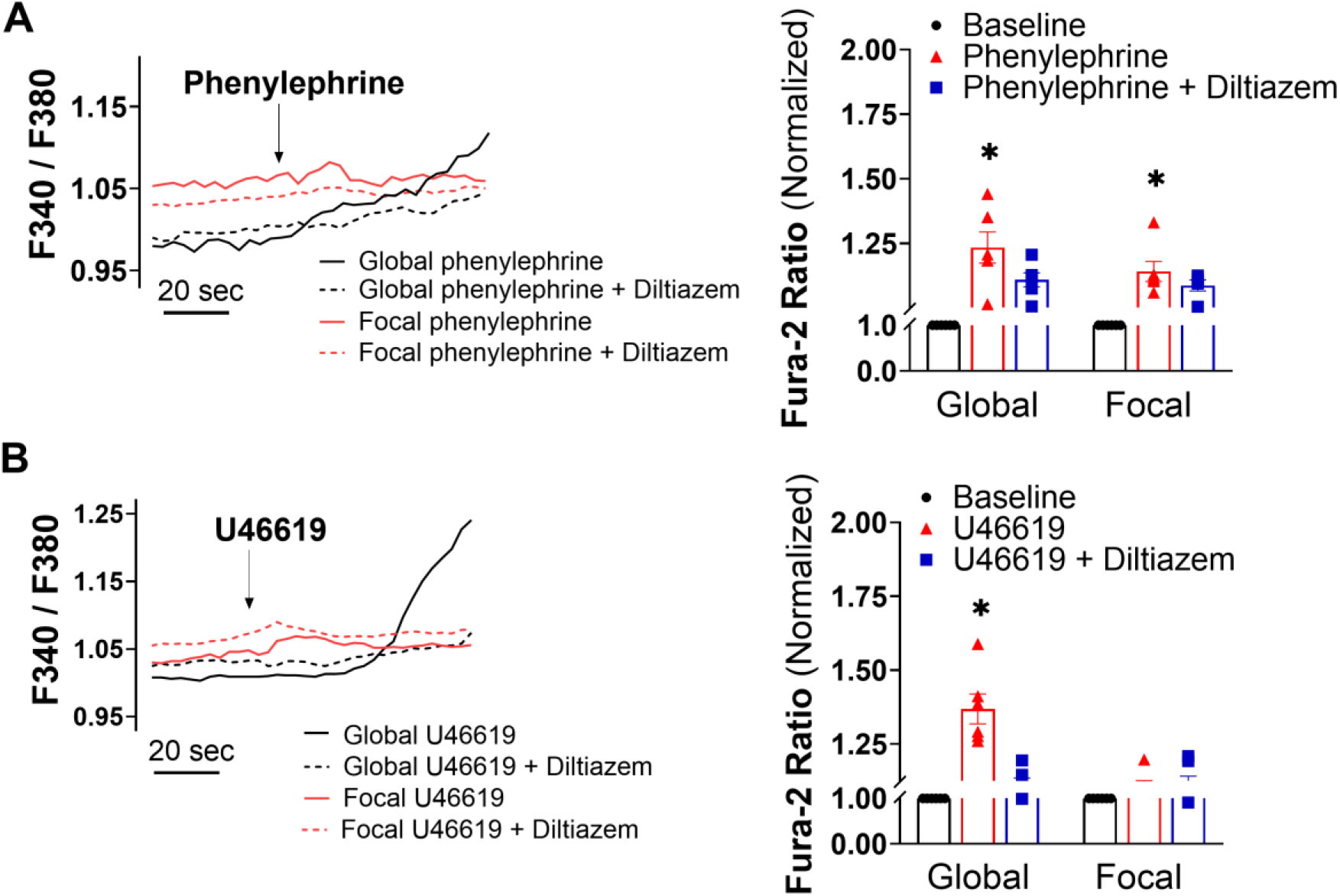
Effects of focal and global agonist application on intracellular [Ca^2+^]. Representative traces and summary data show the cytosolic Ca^2+^ changes (F340/F380) in response to focal and global phenylephrine **(A)** and U46619 **(B)** with and without diltiazem (30 μm, L-type channel blocker). Summative cytosolic Ca^2+^ data are normalized to baseline. N = 6 arteries per group, 1 experiment per mouse. Data are mean ± SEM. * denotes significant decrease from baseline recording, (Paired *t-*test, P-value < 0.05).

## Discussion

This study delved into receptor mediated constriction and the interplay among electro- and pharmacomechanical coupling mechanisms^38^. Work began by constructing concentration response curves to classic G_q/11_ (phenylephrine) or G_q/11_-G_12/13_ (U46619) agonists with and without an L-type Ca^2+^ channel blocker to separate among the coupling processes. Nifedipine diminished and rightward-shifted the concentration-response curves, consistent with electromechanical preceding pharmacomechanical responses, the so-called functional coupling bias. Further analysis of pharmacomechanical coupling revealed that phenylephrine phosphorylated CPI-17, an upstream regulator target of PP1c (MLCP catalytic subunit) where U46619 additionally phosphorylated MYPT1 (MLCP targeting subunit). Complementary tone measures aligned with the biochemistry data, with PKC blockade abolishing the pharmacomechanical response to phenylephrine while that induced by U46619 required both PKC and Rho-kinase inhibition. Subsequent work revealed that the order and dominance of the coupling mechanism “switched” if the number of cells stimulated was restricted though focal rather than global agent application. Clearly then, the coupling mechanisms underlying agonist-induced tone aren’t set in strict relative proportion to one another. Rather, dependent upon the concentration and manner of presentation, agonists trigger a range of vasomotor signatures, functionally biased toward electromechanical or pharmacomechanical coupling. We posit this flexible design is necessary for vascular networks to match blood flow delivery among tissue regions, whose metabolic requirements are disparate and variable with time or other physiological input.

### Functional bias in vascular smooth muscle

Arterial networks are responsible for setting how much and where blood flow is delivered within an organ^1,2^. In peripheral tissues like the gut, arterial tone is defined by the balance between constrictor and dilatory stimuli, the former in response to pressure, sympathetic transmitter release, and paracrine agents derived from endothelium and platelets. Each stimuli works through a G-protein coupled receptor, the resultant constriction dependent on two general mechanisms, an electromechanical process linked to V_M_ and Ca^2+^ influx through L-type Ca^2+^ channels, and pharmacomechanical coupling, a voltage-insensitive pathway tied to MLCP regulation^39^. While studies have long noted that agonists activate both mechanisms, what’s unclear is whether their relative contribution is fixed^5^ or variable, the latter enabled in part the receptor signaling bias that arises as ligand occupancy^40,41^, receptor density^42^ or structural configuration changes^43,44^. It’s in this context that we examined mesenteric contractile responses to two standard agonists, in the absence and presence of nifedipine, an L-type Ca^2+^ channel blocker that separates electro-from pharmacomechanical coupling^45^. The agonists of interest are phenylephrine and U46619, stable analogs of agents released from perivascular nerves and endothelium or platelets, respectively^46,47^. The agonists of interest were phenylephrine and U46619, the former confirmed through use of YM254890 to mediate constriction through G_q/11_ coupled α1-adrenoceptors, the latter via TXA_2_ receptors coupled to G_q/11_ & G_12/13_ (Figure 1)^48^. Irrespective of the agonist and the G-protein coupling, nifedipine attenuated constriction and induced a rightward shift in the concentration response curve (Figure 2). This shift is consistent with a functional signaling bias, with electromechanical coupling preceding but giving way to pharmacomechanical coupling as noted in Figure 2A-B, right. Thus in contrast to coupling mechanisms presumptively activated in fixed relative proportion, our work indicates they are engaged in a defined sequential order^45^. Ensuing analysis also revealed that phenylephrine induced quicker temporal engagement of electromechanical coupling than pharmacomechanical (Figure 2C & 2D). A similar trend wasn’t observed with U46619, a difference difficult to resolve but likely tied to the TXA_2_ receptor’s more complex G-protein coupling arrangement. Note, control experiments confirmed that qualitatively similar findings were obtained in mesenteric arteries lacking Ca_V_3.1, a T-type Ca^2+^ channel with a limited role in tone development (Figure 3)^34^.

### Foundation of pharmacomechanical coupling

While electromechanical coupling, with its linkage to blood flow regulation, has been studied in depth, pharmacomechanical coupling has received decidedly less attention due to noted difficulties in conducting classic western blot analysis on small resistance vessels^12,49^. Classic literature typically ties pharmacomechanical coupling to the regulation of MLCP through its 1) catalytic subunit (PP1c) via CPI-17 phosphorylation and 2) targeting subunit (MYPT1) via T-855 and T-697 phosphorylation (Figure 4A)^50^. In line with this perspective, we performed an amplified three-step western blot approach on mouse mesenteric arteries, with phenylephrine and U46619 application notable for elevating CPI-17 phosphorylation (Figure 4 ) through PKC activation^24^. PKC is a classic downstream target of G_q_ coupled receptors activated through phospholipase C and the production of diacylglycerol and IP_3_; the latter drives Ca^2+^ release from the sarcoplasmic reticulum^51^. Interestingly, subsequent work revealed that while phenylephrine had no effect on MYPT1, U46619 enhanced phosphorylation at sites T-855 and T-697, which are under Rho-kinase control. Rho-kinase is a downstream target of G_12/13_ coupled receptors activated by RhoA, a small GTPase under the regulatory control of guanine exchange factors^52^. Consistent with these biochemical observations, functional work reveals that Calphostin C, a broad-spectrum PKC antagonist eliminated the ability of phenylephrine to induce nifedipine-insensitive constriction (Figure 5). In contrast, PKC and Rho-kinase inhibition were both required to abolish the nifedipine-insensitive constriction by U46619, an agonist that mediates its effects through TXA_2_ receptors that are G_q/11_ and G_12/13_ coupled (Figure 1B)^53^. The variability in the phosphorylation pattern highlights the interplay amongst agonists, signaling pathways and the molecular underpinnings of pharmacomechanical coupling. Final note, while MYTP1 is a recognized Rho-kinase target, its effects on pharmacomechanical coupling may also involve the modulation of actin stress fibers, a worthy subject for future study^54,55,56^.

### Altering functional bias of resistance arteries

Our initial work showed that coupling mechanisms are sequentially, rather than concurrently activated, with electromechanical dominating at low agonist concentrations and pharmacomechanical activating in accordance with dosage. A query that logically follows is whether the functional bias towards electromechanical coupling, presumptively enabled receptor signaling bias, can switch to full pharmacomechanical dominance. Consider in detail the computer model in Figure 6, in which a small depolarizing current is injected into a variable number of smooth muscle cells within the virtual artery. When all are stimulated, we observed a pronounced depolarization of 12.5 mV, a response that starkly contrasts the <2.0 mV depolarization noted when the number of stimulated smooth muscle cells is reduced to 1/8 of total. The latter finding is an intriguing but predictable change that arises when charge from a small number of stimulated cells disperses within the greater mass of unstimulated cells. It suggests that pharmacomechanical coupling is likely to be the dominant contractile mechanism in scenarios where agents are presented focally and discretely. This conceptual insight aligns well with experimentation which shows that focal phenylephrine or U46619 application elicits discrete constrictions unaffected by nifedipine, an agent that separates amongst electromechanical and pharmacomechanical coupling (Figure 7). PKC appears to play a prominent role in mediating these discrete events, as responses were markedly attenuated in the presence of Calphostin C. Note that in subsequent work, we confirmed the absence of arterial depolarization to focal agonist application (Figure 8) while global superfusion-induced robust depolarization (∼10-12 mV). This shift to pharmacomechanical dominance was also evident in our [Ca^2+^]_i_ measures; focal application elicited a modest, nifedipine-insensitive rise whereas the large change induced by global application was abolished by this L-type Ca^2+^ channel blocker (Figure 9). While the mechanisms driving the voltage-insensitive [Ca^2+^]_i_ responses are unclear, past studies have implied a role for: 1) receptor operated, transient receptor potential channels; or 2) Ca^2+^ waves, asynchronous events that originate from the sarcoplasmic reticulum^6,8,57^.

### Physiological significance

The foundation of excitation-contraction coupling is decidedly more complex for vascular tissue than for cardiac or skeletal muscle. In addition to membrane potential driving Ca^2+^ influx (electromechanical), a nonelectrical (pharmacomechanical) component tied to MLCP regulation must be carefully weighted and assigned a physiological role. The latter is difficult to ascertain if one adopts the traditional view that coupling mechanisms are set in fixed relative proportion, as both would presumptively support the same contractile behaviours^58^. Greater clarity, however, emerges as one considers functional bias and the ability of agonists to elicit a diversity of vasomotor “signatures”. First consider vasoactive agents circulating in blood or released from perivascular nerves at low concentration, a scenario where the ensuing vasomotor responses would be biased towards electromechanical coupling. The charge generated in support of depolarization would readily integrate among connected cells, across multiple arterial segments, a scenario ideal for setting base blood flow magnitude across a network. With stronger stimulation and further elevation of concentration, electrical control would cede to pharmacomechanical coupling, with a portion of tone development being resistant to arterial V_M_ modulation. In this scenario, hyperpolarizing responses, due to feedback loops or endothelial activation, would lose their ability to fully dilate an artery. Moderation of pharmacomechanical control would require an alternative nonelectrical mechanism, perhaps one tied to nitric oxide and protein kinase G, which could conceivably limit MLCP inhibition by interfering with MYPT1 phosphorylation^57,58,59^. Note, if agent concentration was high and focal enough, the result of local varicosity release, intrinsic tissue production or injury, small arterial segments would expectantly operate independent of the broader network, to tune blood flow distribution discretely and markedly to defined regions^60^. The idea of local pharmacomechanical control should be carefully weighed in context to arterial vasospasm, a disease state notably insensitive to dihydropyridines^61^. Work herein indicates a more effective means of ameliorating this deleterious state would be to therapeutically target signaling proteins (e.g. PKC or Rho-kinase) within non-electrical (i.e. pharmacomechanical) coupling pathways^62,63,64^.

## Conclusion

This study presents three key findings, the first being that constrictor agonists working through the α1-adenoreceptor or TXA_2_ receptors elicit functionally biased responses, with electromechanical preceding, but giving way to, pharmacomechanical coupling as concentration rises. Second, our work reveals a key role for PKC and Rho-kinase in pharmacomechanical coupling, the former enhancing CPI-17 phosphorylation (regulator of the catalytic subunit PP1c), and the latter MYPT1 phosphorylation (the myosin phosphatase targeting subunit). Lastly, we show that the functional bias toward electromechanical coupling can switch to pharmacomechanical dominance by simply changing how agents are presented to arteries. We conclude that agonists elicit a dynamic range of vasomotor signatures, each presumptively important to the real time control of arterial networks and setting blood flow delivery. Concepts herein also provide new insight on the contractile foundation of arterial vasospasm, with its deleterious knock-on effects.

## Acknowledgments

The authors would like to thank all the members of our research group who contributed to this study. Special thanks are extended to Dr. Peter Chidiac, Suzanne Brett Welsh, and Naman Arora. Schematic illustrations were created with BioRender.com under Dr. Galina Yu. Mironova license.

## Supplemental Material

Figure S4B-D

Data Set

Major Resources Table

## Non-standard Abbreviations and Acronyms

Ca^2+^-CAM: Ca^2+^-calmodulin
CPI-17: Myosin phosphatase inhibitory protein of 17 kDa
EC: Endothelial cell
GPCR: G protein-coupled receptors
MYPT1: Myosin phosphatase target subunit 1
MLC: Myosin light chain 20 kDa
MLCK: Myosin light chain kinase
MLCP: Myosin light chain phosphatase
PKC: Protein kinase C
PP1cδ: Myosin phosphatase catalytic subunit
SMC: Smooth muscle cell
TXA_2_: Thromboxane A_2_
T-855: Threonine-855
T-697: Threonine-697
V_M_: Membrane potential
[Ca^2+^]_i_: Intracellular Ca^2+^ concentration

## References

1. Liu Z, Khalil RA. Evolving mechanisms of vascular smooth muscle contraction highlight key targets in vascular disease. Biochem Pharmacol. 2018; 153:91–122 doi:10.1016/j.bcp. 2018.02.012

2. Cai C, Fordsmann JC, Jensen SH, Gesslein B, Lønstrup M, Hald BO, Zambach SA, Brodin B, Lauritzen MJ. Stimulation-induced increases in cerebral blood flow and local capillary vasoconstriction depend on conducted vascular responses. Proc Natl Acad Sci U S A. 2018; 115(25):E5796–E5804. doi:10.1073/pnas.1707702115

3. Kauffenstein G, Laher I, Matrougui K, Guérineau NC, Henrion D. Emerging role of G protein-coupled receptors in microvascular myogenic tone. Cardiovasc Res. 2012; 95(2):223–232. doi:10.1093/cvr/cvs152

4. Pierce KL, Premont RT, Lefkowitz RJ. Seven-transmembrane receptors. Nat Rev Mol Cell Biol. 2002; 3(9):639–650. doi:10.1038/nrm908

5. Casteels R. Electro- and pharmacomechanical coupling in vascular smooth muscle. Chest. 1980; 78:150–156. doi:10.1378/chest.78.1_Supplement.150

6. Kuo KH, Dai J, Seow CY, Lee CH, Van Breemen C. Relationship between asynchronous Ca^2+^ waves and force development in intact smooth muscle bundles of the porcine trachea. Am J Physiol - Lung Cell Mol Physiol. 2003; 285(6):L1345–53. doi:10.1152/ajplung.00043.2003

7. Somlyo AP, Somlyo AV. Smooth muscle: Excitation-contraction coupling, contractile regulation, and the cross-bridge cycle. Alcohol Clin Exp Res. 1994; 18(1). doi:10.1111/j.1530-0277.1994.tb00893.x

8. Longden TA, Hill-Eubanks DC, Nelson MT. Ion channel networks in the control of cerebral blood flow. J Cereb Blood Flow Metab. 2016; 36(3):492–512. doi:10.1177/0271678X15616138

9. Cole WC, Welsh DG. Role of myosin light chain kinase and myosin light chain phosphatase in the resistance arterial myogenic response to intravascular pressure. Arch Biochem Biophys. 2011; 510(2):160–173. doi:10.1016/j.abb.2011.02.024

10. Moreno-Domínguez A, Colinas O, El-Yazbi A, Walsh EJ, Hill MA, Walsh MP, Cole WC. Ca^2+^ sensitization due to myosin light chain phosphatase inhibition and cytoskeletal reorganization in the myogenic response of skeletal muscle resistance arteries. J Physiol. 2013; 591(5):1235–1250. doi:10.1113/jphysiol.2012.243576

11. Kitazawa T, Eto M, Woodsome TP, Khalequzzaman M. Phosphorylation of the myosin phosphatase targeting subunit and CPI-17 during Ca^2+^ sensitization in rabbit smooth muscle. J Physiol. 2003; 546(3):879–889. doi:10.1113/jphysiol.2002.029306

12. Itoh T. Review: Pharmacomechanical coupling in vascular smooth muscle cells - An overview -. Jpn J Pharmacol. 1991; 55(1):1–9. doi:10.1016/s0021-5198(19)39971-8

13. Segal SS. Cell-to-cell communication coordinates blood flow control. Hypertension. 1994; 23(6):1113–1120. doi:10.1161/01.HYP.23.6.1113

14. Diep HK, Vigmond EJ, Segal SS, Welsh DG. Defining electrical communication in skeletal muscle resistance arteries: A computational approach. J Physiol. 2005; 568(1):267–281. doi:10.1113/jphysiol.2005.090233

15. Kapela A, Bezerianos A, Tsoukias NM. A mathematical model of vasoreactivity in rat mesenteric arterioles: I. Myoendothelial communication. Microcirculation. 2009; 16(8):694–713. doi:10.3109/10739680903177539

16. Chidiac P. RGS proteins destroy spare receptors: Effects of GPCR-interacting proteins and signal deamplification on measurements of GPCR agonist potency. Methods. 2016; 92:87–93. doi:10.1016/J.YMETH.2015.08.011

17. Boivin GP, Bottomley MA, Dudley ES, Schiml PA, Wyatt CN, Grobe N. Physiological, behavioral, and histological responses of male C57BL/6N mice to different CO_2_ chamber replacement rates. J Am Assoc Lab Anim Sci. 2016; 55(4):451–461.

18. Wenceslau CF, McCarthy CG, Earley S, England SK, Filosa JA, Goulopoulou S, Gutterman DD, Isakson BE, Kanagy NL, Martinez-Lemus LA, et al. Guidelines for the measurement of vascular function and structure in isolated arteries and veins. Am J Physiol - Hear Circ Physiol. 2021; 321(1):H77–H111. doi:10.1152/ajpheart.01021.2020

19. Kim D, Song I, Keum S, Lee T, Jeong M, Kim S, McEnery M, Shin H. Lack of the burst firing of thalamocortical relay neurons and resistance to absence seizures in mice lacking α1G T-type Ca^2+^ channels. Neuron. 2001;31(1):35–45. doi:10.1016/S0896-6273(01)00343-9

20. Knot HJ, Nelson MT. Regulation of membrane potential and diameter by voltage-dependent K^+^ channels in rabbit myogenic cerebral arteries. Am J Physiol. 1995; 269(1 Pt 2):H348-55. doi: 10.1152/ajpheart.1995.269.1.H348.

21. Nunes KP, Webb RC. New insights into RhoA/Rho-kinase signaling: a key regulator of vascular contraction. Small GTPases. 2021; 12(5-6):458–469. doi:10.1080/21541248.2020.1822721

22. Norton CE, Jernigan NL, Walker BR, Resta TC. Membrane depolarization is required for pressure-dependent pulmonary arterial tone but not enhanced vasoconstriction to endothelin- 1 following chronic hypoxia. Pulm Circ. 2020; 10(4). doi:10.1177/2045894020973559

23. Takeya K, Loutzenhiser K, Shiraishi M, Loutzenhiser R, Walsh MP. A highly sensitive technique to measure myosin regulatory light chain phosphorylation: The first quantification in renal arterioles. Am J Physiol - Ren Physiol. 2008; 294(6):1487–1492. doi:10.1152/ajprenal.00060.2008

24. Johnson RP, El-yazbi AF, Takeya K, Walsh EJ, Walsh MP, Cole WC. Ca^2+^ sensitization via phosphorylation of myosin phosphatase targeting subunit at threonine-855 by Rho kinase contributes to the arterial myogenic response. J Physiol. 2009; 587(11):2537–2553. doi:10.1113/jphysiol.2008.168252

25. Hald BO, Welsh DG. Conceptualizing conduction as a pliant vasomotor response: impact of Ca^2+^ fluxes and Ca^2+^ sensitization. Am J Physiol - Hear Circ Physiol. 2020; 319(6):H1290–H1301. doi:10.1152/ajpheart.00286.2020

26. Welsh DG, Segal SS. Endothelial and smooth muscle cell conduction in arterioles controlling blood flow. Am J Physiol. 1998; 274(1):H178–86. doi: 10.1152/ajpheart.1998.274.1.H178

27. Tran CHT, Vigmond EJ, Plane F, Welsh DG. Mechanistic basis of differential conduction in skeletal muscle arteries. J Physiol. 2009; 587(6):1301–1318. doi:10.1113/jphysiol.2008.166017

28. Kharche SR, Mironova GY, Goldman D, McIntyre CW, Welsh DG. Sensitivity Analysis of a smooth muscle cell electrophysiological model. Lect Notes Comput Sci. 2021;12738 LNCS:540-550. doi:10.1007/978-3-030-78710-3_52/FIGURES/5

29. Hindmarsh AC, Brown PN, Grant KE, Lee, Steven L, Serban R, Shumaker DE, Woodward CS. SUNDIALS: Suite of Nonlinear and Differential/Algebraic Equation Solvers. ACM Transactions on Mathematical Software. 2005; 31(3): 363–396 10.1145/1089014.1089020

30. Kharche S, Adeniran I, Stott J, Law P, Boyett, MR, Hanco JC, Zhang H. Pro-arrhythmogenic effects of the S140G KCNQ1 mutation in human atrial fibrillation - insights from modelling. J Physiol. 2012; 590(18):4501–4514. doi:10.1113/JPHYSIOL.2012.229146

31. Nishimura A, Kitano K, Takasaki J, Taniguchi M, Mizuno N, Tago K, Hakoshima T, Itoh H. Structural basis for the specific inhibition of heterotrimeric Gq protein by a small molecule. Proc Natl Acad Sci USA. 2010; 107(31):13666–13671. doi:10.1073/pnas.1003553107

32. Takasaki J, Saito T, Taniguchi M, Kawasaki T, Moritani Y, Hayashi K, Kobori M. A novel Gαq/11-selective inhibitor. J Biol Chem. Published online 2004; 279(46) 47438–47445 doi:10.1074/jbc.M408846200

33. Harraz OF, Visser F, Brett SE, Goldman D, Zechariah A, Hashad AM, Menon BK, Watson T, Starreveld Y, Welsh DG. CaV1.2/CaV3.x channels mediate divergent vasomotor responses in human cerebral arteries. J Gen Physiol. 2015; 145(5):405–418. doi:10.1085/jgp.201511361

34. El-Lakany MA, Haghbin N, Arora N, Hashad AM, Mironova GY, Sancho M, Gros R, Welsh DG. CaV3.1 channels facilitate calcium wave generation and myogenic tone development in mouse mesenteric arteries. Sci Rep. 2023; 13(1):1–16. doi:10.1038/s41598-023-47715-3

35. Takashima S. Phosphorylation of myosin regulatory light chain by myosin light chain kinase, and muscle contraction. Circ J. 2009; 73(2):208–213. doi:10.1253/circj.CJ-08-1041

36. Abd-Elrahman KS, Colinas O, Walsh EJ, Zhu HL, Campbell CM, Walsh MP, Cole WC. Abnormal myosin phosphatase targeting subunit 1 phosphorylation and actin polymerization contribute to impaired myogenic regulation of cerebral arterial diameter in the type 2 diabetic Goto-Kakizaki rat. J Cereb Blood Flow Metab. 2017; 37(1) doi:10.1177/0271678X15622463

37. Jiang RS, Zhang L, Yang H, Zhou MY, Deng CY, Wu W. Signaling pathway of U46619- induced vascular smooth muscle contraction in mouse coronary artery. Clin Exp Pharmacol Physiol. 2021; 48(7):996–1006. doi:10.1111/1440-1681.13502

38. Somlyo Be AP, Somlyo A V. Signal transduction and regulation in smooth muscle: problems and progress. Rev Physiol Biochem Pharmacol. 1994; 134:1–6. doi: 10.1007/3-540-64753-8_1.

39. Wilson DP, Susnjar M, Kiss E, Sutherland C, Walsh MP. Thromboxane A2-induced contraction of rat caudal arterial smooth muscle involves activation of Ca^2+^ entry and Ca^2+^ sensitization: Rho-associated kinase-mediated phosphorylation of MYPT1 at Thr-855, but not Thr-697. Biochem J. 2005; 389(Pt3):763. doi:10.1042/BJ20050237

40. Onaran HO, Costa T. Conceptual and experimental issues in biased agonism. Cell Signal. 2021; 82. doi:10.1016/j.cellsig.2021.10995

41. Smith JS, Lefkowitz RJ, Rajagopal S. Biased signaling: from simple switches to allosteric microprocessors. Nat Publ Gr. 2018; 17. doi:10.1038/nrd.2017.229

42. Li A, Liu S, Huang R, Ahn S, Lefkowitz RJ. Loss of biased signaling at a G protein coupled receptor in overexpressed systems. PLoS One. 2023; 18(3):1–15. doi:10.1371/journal.pone.0283477

43. Franco R, Aguinaga D, Jiménez J, Lillo J, Martínez-Pinilla E, Navarro G. Biased receptor functionality versus biased agonism in G-protein-coupled receptors. Biomol Concepts. 2018;9(1):143–154. doi:10.1515/bmc-2018-0013

44. Navarro G, Cordomí A, Zelman-Femiak M, et al. Quaternary structure of a G-protein- coupled receptor heterotetramer in complex with Gi and Gs. BMC Biol. 2016; 14(1):1–12. doi:10.1186/s12915-016-0247-4

45. Kobayashi S, Gong MC, Somlyo A V., Somlyo AP. Ca^2+^ channel blockers distinguish between G protein-coupled pharmacomechanical Ca^2+^ release and Ca^2+^ sensitization. Am J Physiol - Cell Physiol. 1991;260(2 29-2). doi:10.1152/ajpcell.1991.260.2.c364

46. Burnstock G, Ralevic V. New insights into the local regulation of blood flow by perivascular nerves and endothelium. Br J Plast Surg. 1994; 47(8):527–543. doi:10.1016/0007-1226(94)90136-8

47. Schror K. Thromboxane A2 and platelets as mediators of coronary arterial vasoconstriction in myocardial ischaemia. Eur Heart J. 1990; 11:27–34. doi:10.1093/eurheartj/11.suppl_b.27

48. Nakahata N. Thromboxane A2: Physiology/pathophysiology, cellular signal transduction and pharmacology. Pharmacol Ther. 2008; 118(1):18–35 doi:10.1016/j.pharmthera.2008.01.001

49. Knot HJ, Nelson MT. Regulation of arterial diameter and wall [Ca^2+^] in cerebral arteries of rat by membrane potential and intravascular pressure. J Physiol. 1998; 508 (Pt 1):199–209. doi:10.1111/J.1469-7793.1998.199BR.X

50. Ringvold HC, Khalil RA. Protein kinase C as regulator of vascular smooth muscle function and potential target in vascular disorders. In: Advances in Pharmacology. 2017; 78:203–301. doi:10.1016/bs.apha.2016.06.002

51. Neubauer J. Highlighted Topics series: cellular responses to mechanical stress. J Appl Physiol. 2000; 90:1593–1599. doi: 10.1152/jappl.2000.89.4.1253

52. Siehler S. Regulation of RhoGEF proteins by G 12/13-coupled receptors. Br J Pharmacol. 2009; 158(1):41–49. doi:10.1111/j.1476-5381.2009.00121.x

53. Nobe K, Paul RJ. Distinct Pathways of Ca^2+^ sensitization in porcine coronary artery effects of Rho-related kinase and protein kinase C Inhibition on force and intracellular Ca^2+^. Circ Res. 2001; 88(12):1283–90. doi:10.1161/hh1201.092035

54. Nixon GF, Iizuka K, Haystead CM, Haystead TA, Somlyo AP, Somlyo AV. Phosphorylation of caldesmon by mitogen-activated protein kinase with no effect on Ca^2+^ sensitivity in rabbit smooth muscle. J Physiol. 1995; 487(Pt 2):283. doi:10.1113/JPHYSIOL.1995.SP020879

55. Walsh MP, Cole WC. The role of actin filament dynamics in the myogenic response of cerebral resistance arteries. J Cereb Blood Flow Metab. 2013; 33(1):1–12. doi:10.1038/jcbfm.2012.144

56. Moreno-Domínguez A, El-Yazbi AF, Zhu HL, Colinas O, Zhong XZ, Walsh EJ, Cole DM, Kargacin GJ, Walsh MP, Cole WC. Cytoskeletal reorganization evoked by Rho-associated kinase- and protein kinase C-catalyzed phosphorylation of cofilin and heat shock protein 27, respectively, contributes to myogenic constriction of rat cerebral arteries. J Biol Chem. 2014; 289(30):20939–20952. doi:10.1074/JBC.M114.553743

57. El-lakany MA, Welsh DG. TRP channels : a provocative rationalization for local Ca2+ control in arterial tone development. 2024;15: 1–6. doi:10.3389/fphys.2024.1374730

58. Janssen LJ, Lu-Chao H, Netherton S. Excitation-contraction coupling in pulmonary vascular smooth muscle involves tyrosine kinase and Rho kinase. Am J Physiol - Lung Cell Mol Physiol. 2001; 280(4):L666–74. doi:10.1152/ajplung.2001.280.4.l666

59. Jarajapu Y, Knot HJ. Relative contribution of Rho kinase and protein kinase C to myogenic tone in rat cerebral arteries in hypertension. Am J Physiol Heart Circ Physiol. 2005; 289(5):H1917–22. doi: 10.1152/ajpheart.01012.2004

60. Zambach SA, Cai C, Helms HCC, Hald BO, Dong Y, Fordsmann JC, Nielsen RM, Hu J, Lønstrup M, Brodin B, et al. Precapillary sphincters and pericytes at first-order capillaries as key regulators for brain capillary perfusion. Proc Natl Acad Sci U S A. 2021;118(26): e2023749118 doi: 10.1073/pnas.2023749118

61. Athar MK, Levine JM. Treatment options for cerebral vasospasm in aneurysmal subarachnoid hemorrhage. Neurotherapeutics. 2012; 9(1): 37–43 doi:10.1007/s13311-011-0098-1

62. Iwabuchi S, Yokouchi T, Hayashi M, Sato K, Saito N, Hirata Y, Harashina J, Nakayama H, Akahata M, Ito K, et al. Intra-arterial administration of fasudil hydrochloride for vasospasm following subarachnoid haemorrhage: Experience of 90 cases. Acta Neurochir Suppl. 2011; 110(Pt 2):179–181 doi:10.1007/978-3-7091-0356-2_33

63. Bright R, Steinberg GK, Mochly-Rosen D. δPKC mediates microcerebrovascular dysfunction in acute ischemia and in chronic hypertensive stress in vivo. Brain Res. 2007; 1144:146–55. doi:10.1016/j.brainres.2007.01.113

64. Maruhashi T, Higashi Y. An overview of pharmacotherapy for cerebral vasospasm and delayed cerebral ischemia after subarachnoid hemorrhage. Expert Opin Pharmacother. 2021; 22(12):1601–1614. doi:10.1080/14656566.2021.1912013

